# GOLDEN: Gene expression and allele-informed decomposition for genetic heterogeneity in spatial transcriptomic data

**DOI:** 10.64898/2026.10.18.759920

**Authors:** Sangjeong Lee, Tae Lyun Ko, Hyoung-Sam Heo, Junguee Lee, Sung Ki Lee, Hyojung Paik

## Abstract

**Motivation:** Spatial transcriptomics (ST) enables exploration of cellular interactions, including immune responses. However, current methods rely on pathologists, with limitations in addressing genetic heterogeneity, where niches harboring nonclonal cells, including placental or allograft tissues, are present. Deciphering these interindividual mixtures remains challenging.

**Results:** For deciphering interindividual heterogeneity in ST, we developed GOLDEN (Gene expressiOn and aLlele-informed DEcomposition for geNetic heterogeneity), which involves an allele-aware multitask network that encodes spot-level expressions into shared latent representations. Incorporating allele features, dual heads classify spot identity (target, nontarget, or mixed) and estimate target fractions. We utilized synthetic ST data with spot-level ground truth using semiallogeneic single-cell profiles. For these synthetic benchmarks, GOLDEN achieved reliable deconvolution, with high correlation (Pearson’s r = 0.977) and low error (mean absolute error (MAE) = 0.045) in fraction estimation and consistent predictions across real-world STs. These predictions were genetically validated. Using GOLDEN, resolving spot-level genetic heterogeneity is feasible.

**Availability and implementation:** GOLDEN and the simulation code for ST are available at https://github.com/othertics/GOLDEN.

## 1. Introduction

The spatial information of cells represents the fundamental physical foundation of the structure and function of multicellular organisms, including humans. During development, the precise spatial arrangement of cells ensures the appropriate differentiation of complex tissues and organs and plays a critical role in maintaining normal growth and physiological homeostasis [1]. Furthermore, even in fully developed organisms, such spatial information serves as an indicator of microenvironmental alterations, including cell–cell communication and cellular-level responses to environmental change [2]. Capturing these spatial patterns becomes even more paramount under pathological conditions, such as in the case of diseases or infections. For instance, spatial transcriptomics (ST) analyses of human placental tissues infected with SARS-CoV-2 successfully elucidated specific immunological niches that mediate severe inflammatory responses [3]. Consequently, insights into ST are indispensable not only for uncovering the fundamental principles of multicellular biology but also for deciphering the underlying mechanisms of immune responses [4] and complex disease etiologies, including the tumor microenvironment [5].

To capture this critical spatial information, various sequencing- and imaging-based approaches have recently been introduced, enabling unbiased whole-transcriptome profiling while preserving the tissue’s intrinsic histological architecture [6]. In the case of a sequencing-based approach, ST involves the capture and sequencing of RNAs within localized grid regions across consecutive tissue sections, spanning a spectrum of spatial resolutions ranging from tens of micrometers to the subcellular level owing to recent technological advancements (e.g., Visium = 55 μm in diameter, Visium HD = 2 μm) [7]. However, these high-dimensional spatial data are inevitably associated with continuous mixed signals, where transcriptomic inputs from multiple heterogeneous cells are conflated within a single spot. To overcome this technological bottleneck and effectively interpret these convoluted signals, a variety of cell-type deconvolution algorithms have been proposed [8-10]. By leveraging single-cell RNA sequencing datasets as reference templates, these algorithms decompose pooled transcriptomic profiles into statistical proportions of individual cell lineages. By remapping known cellular phenotypes onto a continuous spatial grid, traditional statistics-based algorithms have contributed significantly to delineating the spatial boundaries of distinct cell lineages.

More recently, various artificial intelligence and deep learning-based frameworks have been proposed to enhance the analytical efficiency of ST data [11-16]. For example, in architectures such as DSTG [17], graph neural networks (GNNs) are integrated to capture complex, nonlinear spatial dependencies among neighboring cells, whereas in frameworks such as Cardamom [18], deep multilayer perceptrons (MLPs) are used to model the stochastic variance of transcriptomic data. By demonstrating superior performance in effectively denoising highly sparse, corrupted spatial data and uncovering latent cellular states, these AI-driven models have substantially broadened the computational biology horizons of spatial transcriptomics. Despite these methodological advancements, existing tools and computational frameworks address mainly the identification of the proportions of cell types, including microglia and T cells [11]. These approaches, however, are not designed to resolve genetic heterogeneity, namely, cases in which cells of distinct genetic origin, such as those found in chimerism and mosaicism, coexist within the same spatial niche. Genetic heterogeneity within the tissue niche refers to a phenomenon whereby cells presenting different genetic variations are dispersed within a given spatial region. For example, a single ST spot at the maternal–fetal interface of the placenta, within localized somatic clonal expansion in tumors, or within transplanted organs inevitably captures a complex mixture of cells derived from disparate genetic backgrounds. Although nonclonal cells exist widely across multicellular organisms [19] and under immunological conditions, resolving genetic heterogeneity has been largely unattempted. We reframe this challenge as follows: rather than estimating cell-type proportions alone, we aim to decipher the genetic origins of cells within each spot.

Taken together, deconvolving spatial genetic mixtures in the absence of explicit, single-cell-level ground truth information remains an unresolved and daunting issue. To address this challenge, we propose GOLDEN (Gene expressiOn and aLlele-informed DEcomposition for genetic heterogeneity), which is a novel allele-aware multitask deep learning architecture explicitly engineered to resolve interindividual genetic mixtures within a continuous tissue space. GOLDEN achieves several conceptual advances. First, instead of using absolute expression magnitudes that are highly sensitive to experimental conditions and sequencing depth, GOLDEN effectively mitigates technical batch effects and sequencing depth biases by transforming raw inputs into binary representations that reflect gene activation states. Furthermore, the model is based on a unified multitask framework that couples categorical spot-state classification with continuous proportion estimation, seamlessly harmonizing discrete genetic boundaries and dense proportional gradients within a single synergistic loss function. Validations on synthetic ST benchmark datasets demonstrate that the GOLDEN achieves sufficient precision (Pearson’s r = 0.977 and MAE = 0.045). Moreover, when applied to unlabeled real human placental ST data, the model successfully generates cell distribution maps that rigorously align with qualitative histological boundaries and orthogonal genotype concordance benchmarks while perfectly preserving macroscopic spatial coherence, as evidenced by the Moran’s I value and neighbor mean absolute error (MAE). By providing a reference-free, allele-aware approach for resolving genetic heterogeneity at the spot level, GOLDEN offers a generalizable framework that is applicable to diverse contexts of nonclonal cell admixture such as maternal–fetal interfaces, allograft rejection, and tumor clonal heterogeneity, thereby extending the analytical scope of spatial transcriptomics beyond conventional cell-type deconvolution.

## 2. Methods

### 2.1. Study approval

The acquisition, processing and storage of human placenta samples were approved by the Konyang University Hospital Institutional Review Board. All the samples were deidentified to protect private information. Clinical characteristics are listed in Supplemental Table 1.

### 2.2. Spatial transcriptomic data generation

Fresh human placental tissues were collected from patients with normal pregnancies and preeclampsia immediately after delivery. Human placenta samples from regions of interest, including decidua and superficial placenta, were prepared as fresh frozen (FF) or fixed frozen (FxF) samples. Blocks were cryosectioned and H&E stained directly on Visium Gene Expression slides (10X Genomics) according to the manufacturer’s standard protocol. The tissues were permeabilized, and the RNA was subjected to spatial transcriptomics library preparation. The tissues were subsequently washed several times with ice-cold PBS (1×; pH 7.4) to remove residual blood and debris. Target regions were dissected using RNase-free instruments and trimmed to fit standard cryomolds. The dissected tissue blocks were embedded in Tissue-Tek OCT compound (Sakura Finetek) and snap-frozen using dry ice. The frozen OCT blocks were stored at −80 °C until sectioning.

Cryosections (6 μm) were cut using a Leica cryotome with the chamber temperature maintained at −20 °C. Representative sections were stained with hematoxylin and eosin (H&E) to verify the presence of the target histological structures. Following histological confirmation, the OCT blocks were shipped on dry ice to Geninus Inc. (Seoul, Korea) for downstream processing. Samples passing quality control (DV200 ≥ 30%) were processed using the 10x Genomics Visium CytAssist Spatial Gene Expression platform with the Visium V4 Slide (FFPE v2 chemistry) and the Visium Human Transcriptome Probe Set.

### 2.3. Spatial transcriptomics, single-cell transcriptomics and genetic variant analyses

Reads of spatial transcriptomics were demultiplexed and aligned to the reference of *Homo sapiens* (GRCh38) using SpaceRanger (v3.1.3). The single-cell transcriptomics sequencing data were aligned and quantified using the Cell Ranger Single-Cell Software Suite (10x Genomics) against the *Homo sapiens* genome (GRCh38). Downstream analyses were conducted using the package Scanpy (version 1.9.1). Variant information was captured at each spot of interest using VarTrix (https://github.com/10xgenomics/vartrix). Although VarTrix is a tool for identifying genetic variant calls in single-cell data on the basis of a set of variants previously defined, we introduced it to capture the allele distribution from each spatial spot identified with Visium. To quantify the genetic heterogeneity of each spot, reads of alternative alleles from each spot were profiled using VarTrix, and the degree of concordance was subsequently calculated against the genotype variance data for fetal cord blood acquired using an Illumina GSA v3 genotype array. In addition, the captured alternative allele count (AAC) for each spot region from synthetic and real-world STs was utilized as an input feature for GOLDEN. All of our ST data, including genotypes, are available for research purposes (GSE341398).

### 2.4. GOLDEN algorithm

#### Framework

In previous studies using single-cell reference data, typical ST deconvolution methods were used to estimate the proportions of multiple cell types within each spot region. However, our approach focuses on characterizing each spot region according to its genetic origin, including target and nontarget cells, and specifically aims to quantitatively predict the target cell (e.g., fetal cell or other genetically distinct cells) proportion within each spot region. In addition, our method is a reference-free approach. To achieve this goal, we designed a multitask learning framework that jointly performs spot identity classification—categorizing spots into target (fetal), nontarget (maternal), or mixed states—and target cell proportion regression. An overview of the proposed GOLDEN framework is provided in Supplementary Figure 1.

#### Preprocessing

Spatial transcriptomics data were represented with a spot-by-gene expression matrix *X* = [*x*_1_, *x*_2_, … *x*_*N*_]^*T*^ ∈ ℝ^*N*×*G*^, where *N* denotes the number of spots and *G* denotes the number of genes. Here, each row *x*_*i*_ ∈ ℝ^*G*^ represents the raw gene expression vector for spot *i*. A total of approximately 38K genes were utilized as input features in this study. Rather than directly utilizing raw gene expression values, we converted the expression profile of each spot into a binarized vector 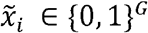 on the basis of expression presence. Specifically, each element of 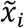 was set to 1 if the corresponding raw value in *x*_*i*_ was greater than 0 and 0 otherwise. This representation emphasizes the presence or absence of gene expression rather than the absolute magnitude of expression. Such a transformation reduces the influence of sequencing depth and library size variability while enabling the model to robustly learn marker gene activation patterns associated with target and nontarget cell compositions.

#### Label Construction

To train the multitask learning framework, we leveraged the predefined cell proportions from synthetic ST datasets as ground-truth labels. For each synthetic spot *i*, the known target cell proportion (e.g., fetal cell fraction) was defined as the continuous regression target, denoted as *f*_*true,i*_. Concurrently, to guide the auxiliary classification task, a discrete three-class label (*y*_*cls,i*_) was constructed by thresholding this ground-truth proportion as follows:

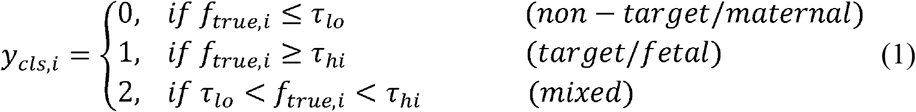

where *τ*_*l o*_ and *τ*_*hi*_ denote the lower and upper thresholds for state determination, respectively. These constructed labels (*f*_*true*_ and *y*_*cls*_) serve strictly as supervised targets during the training phase when synthetic data are utilized. In contrast, during inference based on real-world ST datasets, the model predicts these values directly from the input features without requiring any ground-truth labels.

#### Model Architecture

To learn biologically meaningful spot-level representations, we employed a neural network architecture comprising an MLP-based encoder. For each spatial spot *i*, the binarized gene expression vector 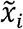 serves as the input to the model. This high-dimensional input is compressed into a lower-dimensional latent representation through a multilayer perceptron (MLP)-based encoder:

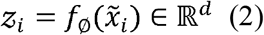

where *f*_Ø_ denotes the encoder network parameterized by weights in set Ø and *z*_*i*_ represents the *d*-dimensional latent embedding vector for spot *i*. Within the encoder, each hidden layer is followed by a Gaussian error linear unit (GELU) activation function and dropout regularization. Layer normalization is exclusively applied to the second hidden block to balance representational capacity and training stability. The resulting latent representation *z*_*i*_ is subsequently provided to both the classification and proportion prediction heads to facilitate multitask learning.

#### Classification Head

The classification head receives the latent representation *z*_*i*_ as the input and predicts a three-dimensional probability vector 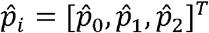 representing three distinct biological states: nontarget (maternal; class 0), target (fetal; class 1), and mixed (class 2). This auxiliary task involves explicitly learning discrete state boundaries that may be difficult to obtain through continuous proportion regression alone, thereby providing structural context to guide the concurrent proportion prediction task.

#### Proportion Head

The proportion head predicts the final target cell fraction 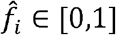 for spot *i* by integrating three complementary information sources: the latent feature vector *z*_*i*_, the alternative allele count (AAC) feature vector *a*_*i*_, and the predicted class probability vector 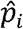 from the classification head. Specifically, these features are concatenated to form a joint representation vector,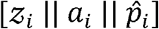. The final prediction is generated through a regression network *g*_ψ_ followed by a sigmoid activation function (σ), ensuring that the predicted fractions naturally remain within the bounded range of [0, 1]:

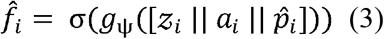

Unlike for discrete state labels, cellular composition (i.e., the fraction of target or non-target cells) is inherently continuous in this case. Therefore, compared with direct prediction from gene expression alone, incorporating both structural state probabilities and AAC-derived genetic evidence enables more robust, biologically informed proportion predictions.

#### Multitask Coupling

A key characteristic of the proposed framework is the explicit coupling of dual tasks, where the proportion head utilizes the predicted class probability vector 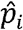 from the classification head as an input feature. Consequently, classification and regression are not optimized independently but are jointly guided through a shared representation. Specifically, the classification task provides coarse-scale structural constraints regarding the biological states (target, nontarget, or mixed), whereas the regression task captures fine-scale, continuous variations within those states. This hierarchical conditioning approach allows the framework to contextualize th e microenvironment state while simultaneously accurately predicting the target cell fraction 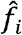, thereby significantly improving both training stability and biological interpretability.

#### Loss Functions

The model is trained using a multitask objective that jointly optimizes classification and proportion prediction tasks to learn shared representation geometries.

#### Classification Loss

To address potential class imbalance issues among the target (fetal), nontarget (maternal), and mixed spots, class-specific weights are integrated into the objective function. Specifically, a multiclass focal loss is used to downweight easy-to-classify samples and focus the model on challenging boundary instances:

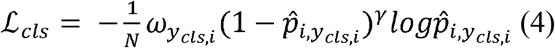

where *y*_*cls,i*_ denotes the ground-truth discrete state label for spot *i*, 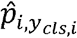 represents the probability predicted by the model for that true class, 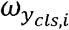, represents the class-specific loss weight, and *γ* represents the focusing parameter.

#### Proportion Regression Loss

The regression objective minimizes the discrepancy between the predicted and true target cell fractions using the mean squared error (MSE):

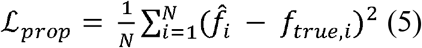

where *f*_*true,i*_ and 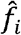 represent the ground-truth and predicted target cell fractions for spot *i*, respectively.

#### Consistency Loss

To enforce strict alignment between the predicted target cell fraction and the discrete state boundaries, we introduced a class-specific constraint loss. This constraint penalizes cases where the continuous proportion prediction contradicts the predicted structural state. Specifically, for a spot assigned to a ground-truth class,*c* ∈ {0, 1, 2} the individual penalty *l*_*i*_ is defined using a hinge-style formulation based on the state-specific thresholds *τ*_*l o*_ and *τ*_*hi*_ :

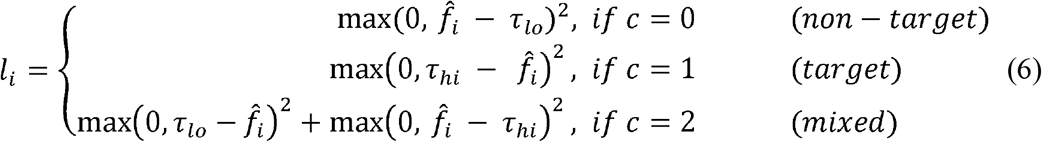

To prevent the consistency objective from being dominated by the majority class within a given training iteration, the penalties are macroaveraged. The individual losses are first averaged within each class and subsequently averaged across all valid classes present in the current mini-batch:

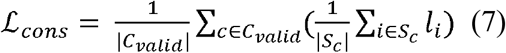

where *S*_*c*_ denotes the set of spots belonging to class *c* in the mini-batch and *C*_*valid*_ = {*c* | |*S*_*c*_| > 0} represents the set of active classes.

#### AAC Loss

For spots with valid alternative allele count (AAC) measurements, we introduce auxiliary AAC consistency loss to anchor the continuous outputs with genetic evidence. First, a predicted microenvironment purity score, *ũ*_*i*_ ∈ [0, 1], is derived from the predicted target fraction 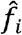:

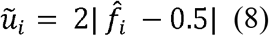

This formulation scales the purity score symmetrically, yielding a value of 0 for perfectly mixed states 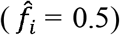 and 1 for entirely pure states 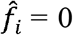. AAC loss is then used to promote alignment between this predicted purity score and the AAC-derived ground-truth purity target *u*_*aac,i*_:

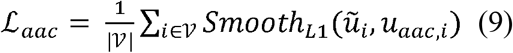

where *V* denotes the set of spots containing valid AAC profiles and *Smooth*_*L*1_ represents the Huber loss function, which is robust against potential sequencing noise in the allele counts. This auxiliary objective successfully incorporates independent genetic evidence into model optimization, preventing overfitting to superficial patterns in simulated expression data.

#### Total Loss

The final multitask training objective is formulated as a weighted linear combination of all primary and regularization losses:

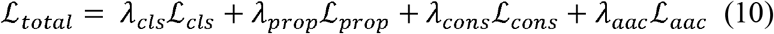

where *λ*_*cls*_, *λ*_*prop*_, *λ*_*cons*_, and *λ*_*aac*_ are hyperparameters that balance the magnitude of each loss component. While the classification and regression objectives serve as the primary driven optimization targets, the consistency and AAC losses function as structural regularizers that ensure biologically coherent predictions.

### 2.5. Generation of synthetic spatial transcriptomics using single-cell transcriptomics

To benchmark our model for spot-level deconvolution of target (fetal) and nontarget (maternal) cell fractions under known ground-truth conditions, we developed a simulation pipeline that synthesizes pseudo-Visium spatial transcriptomics (ST) data from single-cell RNA sequencing (scRNA-seq) data. The pipeline simultaneously generates per-spot cell compositions as ground-truth labels and an aggregated genotype signal (alt_count_sum) derived from VartRix as an orthogonal validation feature. The full pipeline is available at https://github.com/othertics/GOLDEN.

#### Single-cell data preparation for synthetic ST analysis

Raw scRNA-seq data processed by Cell Ranger were loaded, and quality control metrics were computed. To match the per-spot UMI range estimated from the Visium data, each cell was downsampled to a maximum of 2,600 UMI counts using nonreplacement subsampling (scanpy.pp.downsample_counts; replace=False). This target was calibrated so that the summed UMI of cells composited into a single spot (mean ~2–3 cells per spot) fell within the empirically observed Visium range of 5,000–15,000 UMI per spot. Gene identifiers were standardized to Ensembl IDs prior to downstream processing.

#### Genotype signal integration for synthetic ST

To generate an orthogonal genotype-based ground-truth signal for distinguishing fetal from maternal cells, we ran VartRix (scoring-method coverage) against a population-level SNP VCF derived from a public variant database. The number of alternative allele reads summed across all queried SNP loci (alt_count_sum) was computed per cell barcode and merged into the single-cell metadata prior to downsampling.

#### Spatial composition design for synthetic ST

Considering number of spots in real-world ST, a simulated ST slide of 4,000 spots was arranged on a 64 × 64 grid. Maternal cells were assigned to all the spots to model their ubiquitous presence, whereas fetal cells were restricted to a contiguous invasion zone originating from a randomly selected edge position, which is consistent with the known biology of trophoblast invasion. A niche (i.e., fetus–mother mixed spot) was defined as a semicircular region with its center at the grid boundary; the radius was derived analytically from a target spot count sampled from a gamma distribution (mean = 20% of total spots), with area compensation applied to account for the fraction of the circle falling outside the grid. Within the invasion zone, the probability of fetal cell assignment at each spot followed a linear gradient from 1.0 at the zone center to 0.0 at its perimeter. The number of cells per spot was drawn from a Poisson distribution (λ = 2.5), and a minimum of one maternal cell per spot and one fetal cell per invasion-zone spot were enforced to prevent empty assignments.

#### Spot assembly for synthetic ST analysis

For each spot, cells of each type were sampled with replacement from the corresponding single-cell pool, and their gene expression profiles were summed to produce a pseudobulk count vector. Cell types with fewer than 40 cells were excluded from the pool to ensure sampling diversity. Simultaneously, the alt_count_sum values of the sampled cells were aggregated to yield a per-spot genotype signal. The procedure was repeated for independently drawn random seeds to assess simulation variability.

## 3. Results

### 3.1. Benchmark setting based on synthetic and real-world ST data

To train and evaluate GOLDEN with ground truth data, we established synthetic spatial transcriptomic (ST) data from our previous analyses involving placental single-cell RNA-seq (scRNA-seq) with paired genetic evidence from fetal cord blood or maternal blood [20, 21] (Table 1). The source comprised 17 samples spanning three tissue types (maternal–fetal interface (MF), deep placenta (Pla), and endometrium) and diverse clinical conditions, including (normal birth (Norm), preeclamptic preterm birth (PePT), recurrent pregnancy loss during the early trimester (RPL), and normal termination during the early trimester (NormT). Each cell was assigned a maternal (nontarget cell) or fetal (target cell) origin label, as in previous attempts using genetic evidence. Using karyotyping, we excluded one sample from a pregnant women with fetal genetic abnormalities to prevent shortcut learning by the model. These profiles were composited into pseudo-Visium slides designed to mimic the spatial organization of the maternal–fetal interface in the placenta. As described in the Methods section, maternal cells were distributed in a tissue-wide background, and fetal cells were localized to niche-like regions, with three pseudo-Visium slides of randomly sampled cells generated from each of the scRNA-seq datasets. In total, fifty-one pseudo-Visium slides were generated, after which the slides were divided into training (39 slides), validation (6 slides), and test (6 slides) sets. Each spot was assigned an exact target label (fetal cell) proportion as ground truth case together with a nontarget only (maternal-only), target-only (fetal-only), or mixed state label, as well as an alternative allele count (AAC) feature, independent of gene expression.

**Table 1.**
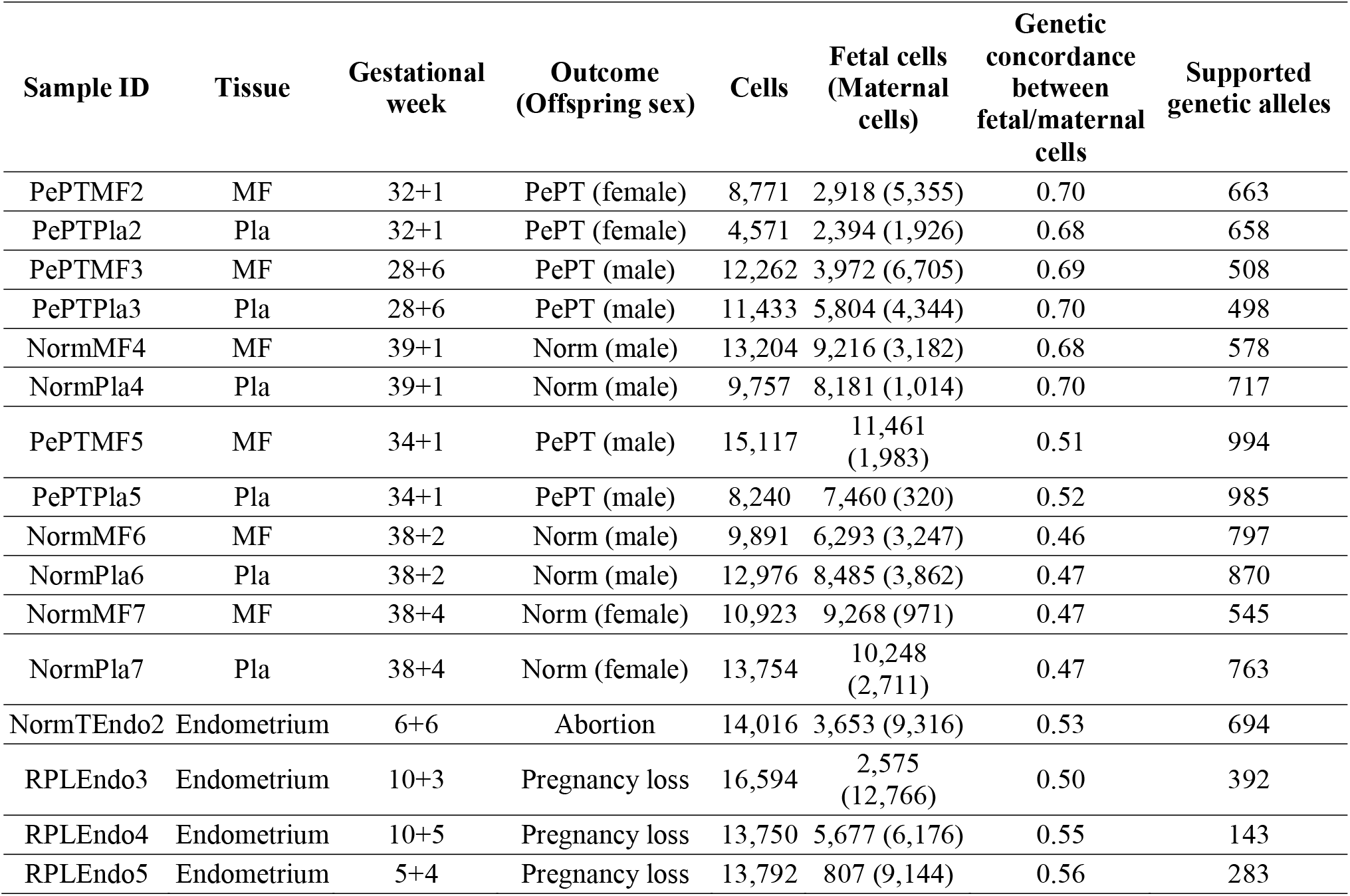
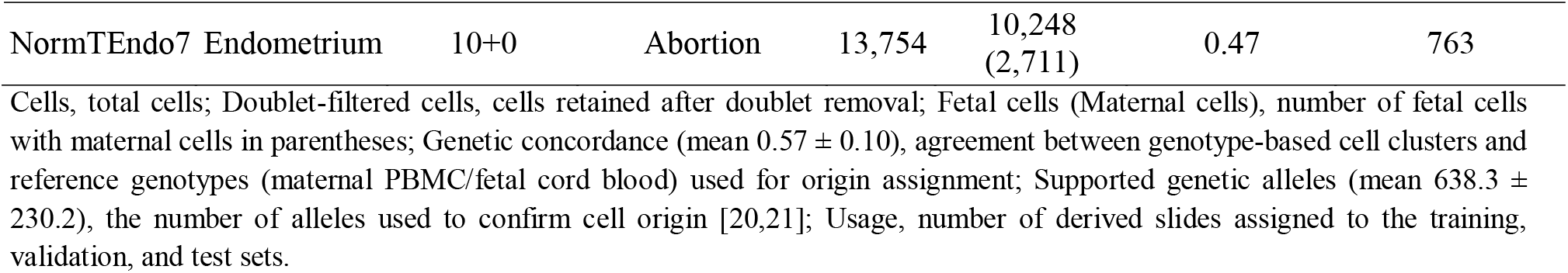
Statistics of scRNA-seq used to generate synthetic ST data.

To evaluate the effects of GOLDEN on real-world ST data, we introduced placental Visium STs from two sources (Supplemental Table 1): internal data from a normal placenta and a preeclamptic placenta and placental ST data from an external cohort of eight individuals [4]. Unlike the synthetic ST data, these real-world ST data lack a per-spot fetal proportion for ground truth referencing. Thus, we calculated the genetic concordance of each spot using genomic variant calls from fetal cells in the corresponding cord blood (details in the Methods section). However, the external data from Arutyunyan et al. did not include genetic variants from either fetal (cord blood) or maternal (maternal blood) cells. Thus, to evaluate the predictions of GOLDEN using the external data, we utilized cell-type annotations from conventional pathology by assigning the cell types of maternal origin, including the decidua and endometrium.

#### Model Performance in Simulation Tests

To objectively evaluate the generalization capability of the proposed GOLDEN model, we performed comprehensive benchmarking based on simulation tests. As detailed previously, the model’s performance was validated using six independent validation datasets that were strictly excluded from the training phase. As illustrated in Figure 1A, the overall performance metrics across the entire test suite yielded an MAE of 0.045, a root mean square error (RMSE) of 0.075, and a Corr. of 0.977, thus demonstrating that GOLDEN robustly estimates the true proportions with high precision. Furthermore, Figure 1B quantitatively delineates the absolute errors across distinct true proportion intervals. The interval-specific MAE ranged from a minimum of 0.031 to a maximum of 0.095. Although a marginal increase in error was observed in the intermediate intervals—reflecting the intrinsic complexity associated with mixed-signal deconvolution—the model nevertheless maintained a remarkably low overall error profile and high predictive stability. The synthetic ST panel in Figure 1C shows a representative comparison between the ground-truth fetal cell proportion map and the inference-based map generated by GOLDEN for the Simulation 615 dataset. GOLDEN reconstructed a spatial distribution nearly identical to that of the original layout, achieving an MAE of 0.033, an RMSE of 0.054, and a Corr. of 0.990. Spatial distribution comparisons between the ground-truth target cell fraction maps and the inference-based maps generated by GOLDEN for the remaining five simulation test samples are provided in Supplementary Figure 2.

**Figure 1.**
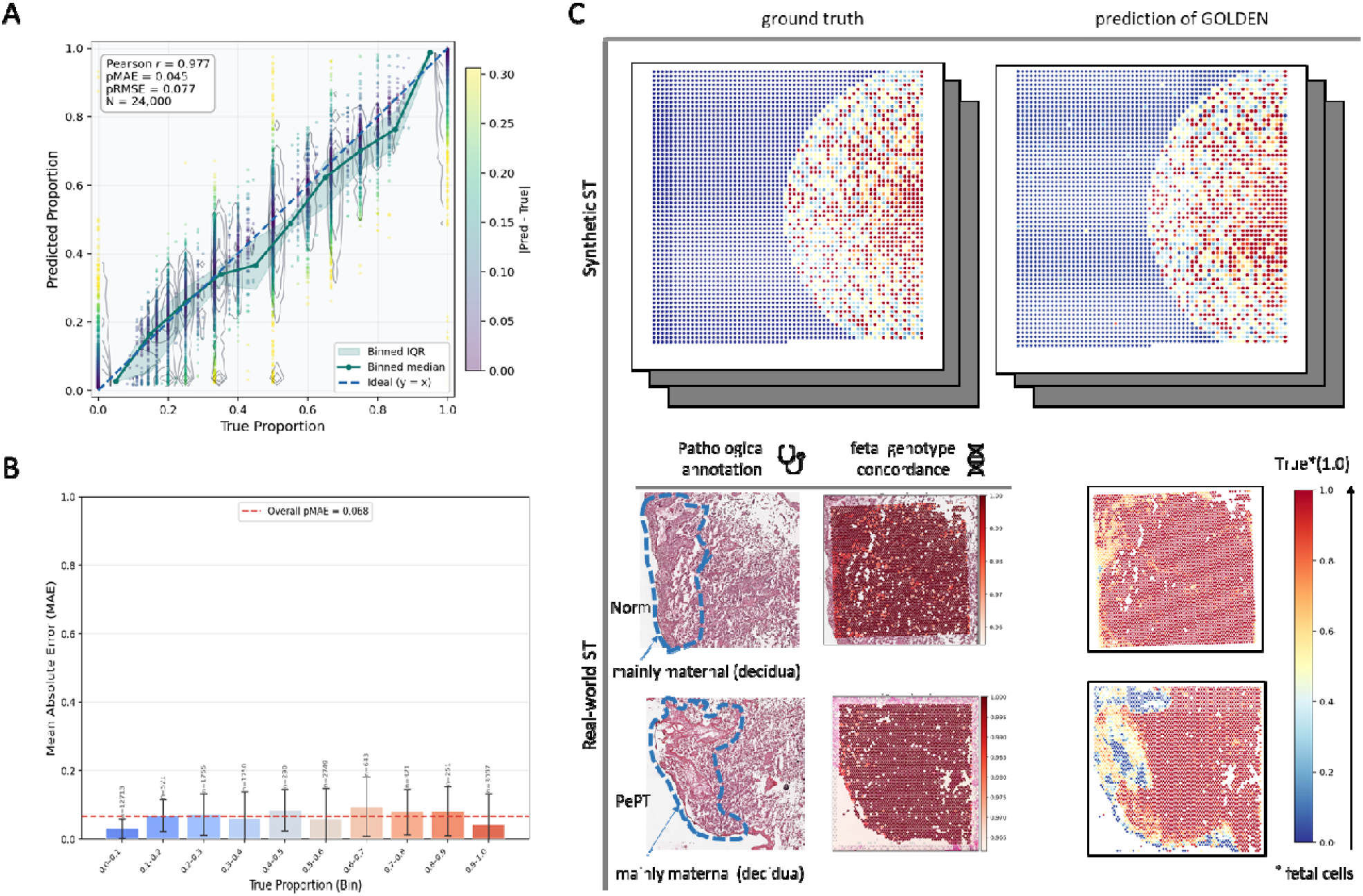
Synthetic ST-based performance evaluation of the proposed model. (A) Regression calibration map illustrating the relationship between the predicted and true fetal proportions, with error shading and calibration ribbons displayed. (B) Error profiling across predicted proportion intervals, demonstrating the variance in absolute proportion errors across the entire prediction spectrum. Across all panels, the proposed model maintains a consistent regression, exhibiting stable calibration and limited error accumulation throughout the proportion range. (C) Synthetic ST: Comparison of the ground-truth fetal cell proportion map and the corresponding inference map generated by GOLDEN for the Simulation 615 dataset. Real-world ST: Spatial proportion inference results obtained with GOLDEN for actual human Norm and PePT samples, presented alongside pathological annotations and fetal genotype concordance information.

#### Model Performance in Real-World Tests

We validated whether the GOLDEN model, trained with simulation data, operates robustly in cases with real-world datasets. Owing to the inherent absence of ground truth information associated with actual tissue data, quantitative performance evaluation is naturally constrained; however, validation can be achieved indirectly using clinicians’ histological annotations of cell types from maternal origin or cord blood genotype concordance metrics. To this end, we analyzed the clinicians’ evaluation results, genotype concordance metrics, and fetal cell proportion maps along with state classification outcomes inferred by GOLDEN, focusing representatively on the Norm and PePT samples (Figure 1C, real-world ST panel). Here, the target and nontarget classes of the model correspond to fetal and maternal origins, respectively. The validation demonstrated that the spatial patterns of maternal and fetal origins inferred by GOLDEN align strongly with the clinicians’ pathological annotations of histological cell types, such as the decidua region for maternal cells. Furthermore, the results of the cord blood genotype validation confirmed that the model’s estimates of allele-informed fractions are in exceptional agreement with known biological evidence (Figure 1C). In addition, we examined predictions of GOLDEN across eight external sample datasets spanning distinct gestational weeks and placental regions (Supplemental Table 1). As depicted in Figure 2A and Supplementary Figure 3, the predictions generated by GOLDEN for these external data align favorably with the cell-type annotations provided by pathologists.

**Figure 2.**
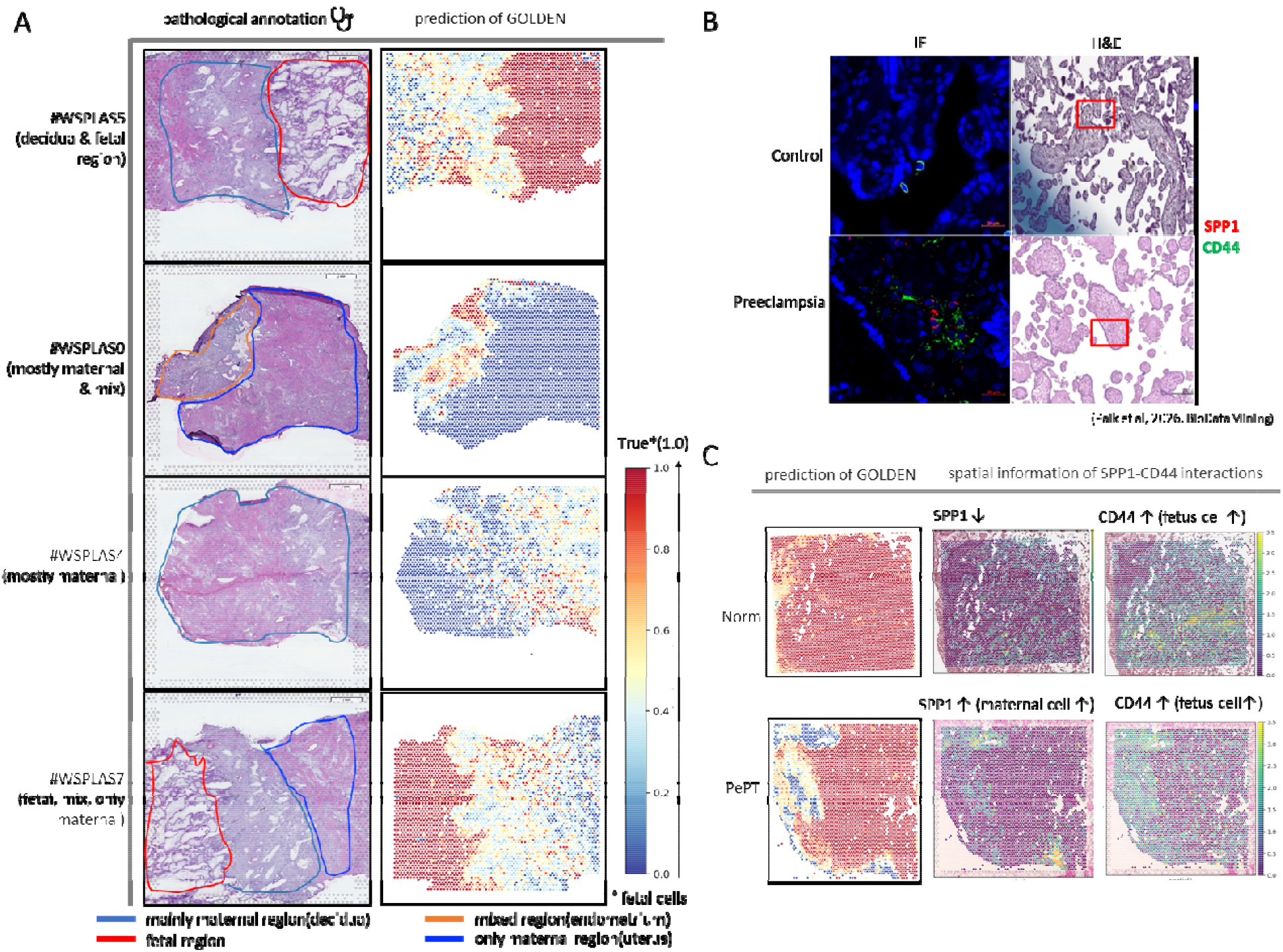
Real-world-based performance evaluation of the proposed model and experimental validation. (A) Comparison of pathological annotations and predictions of GOLDEN using external data (Supplementary Table 1, E-MTAB-12698). (B) Experimental validation of preeclampsia-specific interactions between SPP1 and CD44. Using immunofluorescence (IF) and hematoxylin and eosin (H&E) staining, IF revealed the colocalization of SPP1 (red) and CD44 (green) proteins between cells. H&E staining with a red box indicates immune response activation of cells, including T cells. (C) Predictions of GOLDEN from independent placenta samples under identical disease conditions (e.g., preeclampsia) with confirmation of the genetic origin of cells and matched CD44 and SPP1 expression.

On the basis of the confidence of the predictions of GOLDEN, we further examined whether the predicted genetic heterogeneity of each spot region aligned with the biological findings. Images of colocalization of preeclampsia-associated SPP1-CD44 interactions with unknown cell origin, including fetal or maternal cells, are shown in Figure 2B [20]. Although we previously identified specific interactions in the placenta of preeclampsia patients, the origin of these interactions remains unclear. However, the predictions of GOLDEN revealed proportions of fetal-maternal cells with integrated expression of SPP1 and CD44, confirming that these ligandUreceptor interactions are based on fetal-maternal cells (Figure 2C). Taken together, to the best of our knowledge, GOLDEN is the first tool that can decipher the genetic heterogeneity of spatial regions in ST data. By integrating the inherent transcriptional signatures of ST features for corresponding spatial regions, GOLDEN identifies the niches of semiallogenic cells and related transcriptional signatures.

### 3.2. Ablation study

#### Component Ablation and Trade-offs

To evaluate the contribution of each key component in GOLDEN, we compared the full BASE model against three modified models on the basis of feature ablation: NoCls (no classification head), ExprIn (directly utilizing normalized expression values instead of binarized inputs), and NoAAC (no alternative allele counts) (Figure 3). In the ablation study, we labeled GOLDEN with all features as the BASE model and then compared the modified models on the basis of ablation features. Performance was evaluated across the same six test slides of our synthetic ST data. All the metrics were computed independently at the spot level for each slide and then averaged.

**Figure 3.**
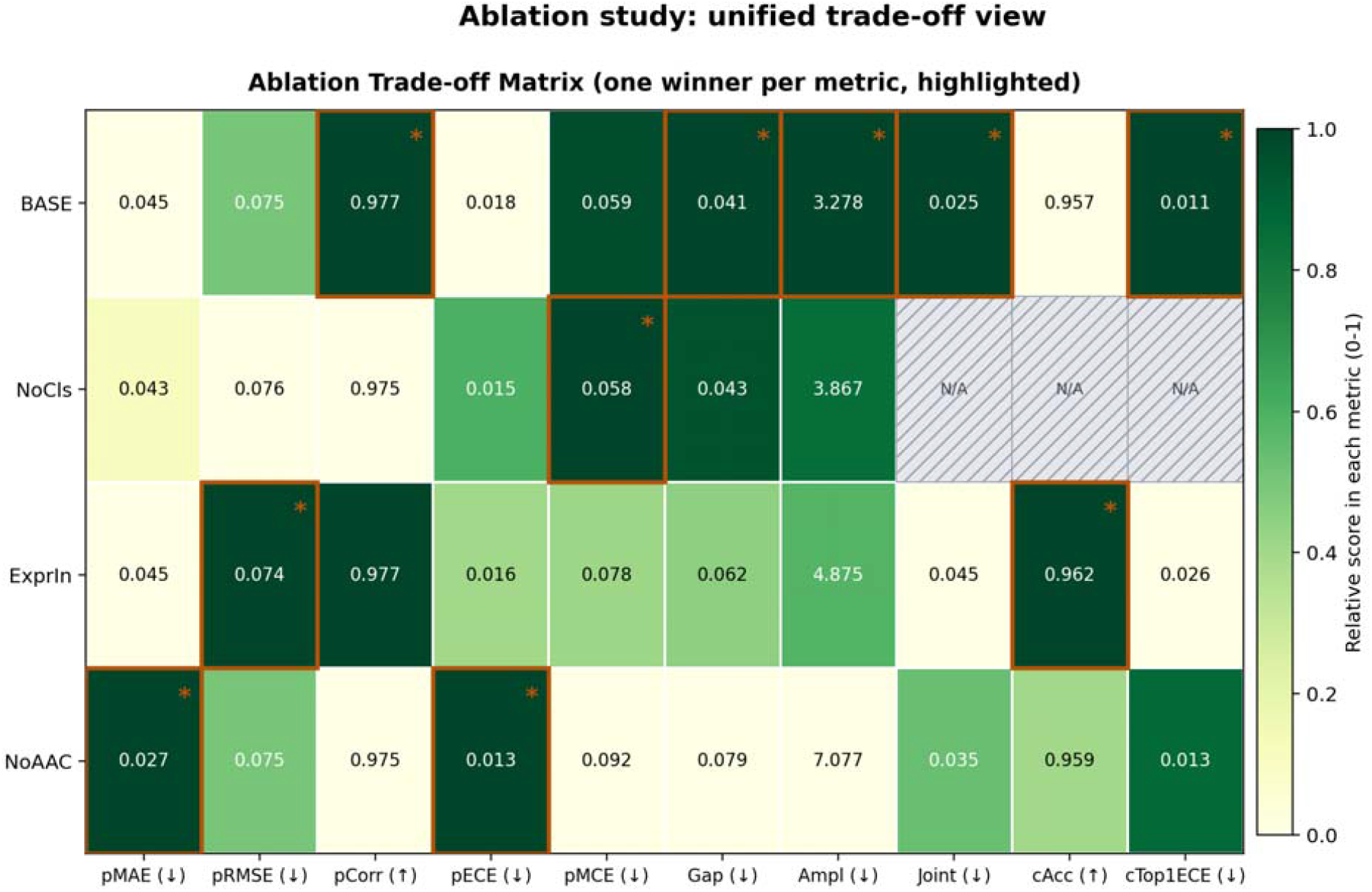
Ablation study-based trade-off matrix across model variants and evaluation metrics. Each row represents a compared model (BASE, NoCls, ExprIn, and NoAAC), while each column denotes regression accuracy, calibration, classification performance, and derived trade-off metrics. The color of each cell indicates the relative performance across that specific column, illustrating the degree of advantage (with higher or lower values being favorable depending on the metric). Asterisks (^*^) and orange borders highlight the top-performing model for each metric.

Given that the proportional prediction of target cells is a continuous regression task, standard classification accuracy cannot be directly applied without the introduction of an arbitrary threshold. To comprehensively evaluate both predictive accuracy and error magnitude, regression performance was evaluated using pMAE (proportion of mean absolute error), pRMSE (proportion of root mean squared error), and pCorr (proportion of Pearson correlation coefficient). Furthermore, to verify the reliability of the model’s confidence and prevent overconfident predictions, calibration quality was evaluated using pECE (proportion of expected calibration error), pMCE (proportion of maximum calibration error), Gap (calibration gap), Ampl (risk amplification), and Joint (joint calibration). For the discrete three-class state classification task, performance was evaluated using cAcc (classification accuracy) and cTop1ECE (classification top-1 expected calibration error) to simultaneously monitor both prediction accuracy and confidence.

NoCls (no classification head model) achieved strong regression accuracy (pMAE = 0.043) and excellent calibration (pECE = 0.015, pMCE = 0.058) performance, but its high pRMSE (0.076) suggested vulnerability to outliers, and its missing head precluded three-class state evaluation. ExprIn (utilizing normalized expression inputs) had the lowest squared error (pRMSE = 0.074) and the highest cAcc (0.962). However, it exhibited poor confidence calibration, marked by an elevated cTop1ECE of 0.026, a high Gap (0.062), and a Joint metric of 0.0450. NoAAC (no alternative allele count model) yielded the lowest pMAE (0.027) and low average calibration error (pECE = 0.013) but displayed extreme tail risk, as evidenced by the highest pMCE (0.092) and severe calibration degradation (Gap = 0.079, Ampl = 7.08).

The full BASE model maintained a well-balanced overall profile (pMAE = 0.045, pRMSE = 0.075, pCorr = 0.977; Figure 3). While its pMAE was slightly higher than that of NoAAC, BASE excelled in terms of worst-case stability and calibration consistency. In summary, while individual compared models showed marginal advantages in terms of some metrics, these gains were accompanied by critical trade-offs. The full BASE model of GOLDEN systematically minimized tail risks—where prediction confidence collapses in challenging subintervals—while maintaining near-peak performance across all dimensions. Consequently, GOLDEN (the full BASE model) was selected as the most robust architecture.

#### Spatial transfer of ablation models for real-world ST data

Although the presented ablation study confirmed the superior performance of the full BASE model of GOLDEN, such results using real-world ST data remain unverified because of the absence of ground truth information. Thus, we quantitatively evaluated the spatial consistency of each model for real-world human placental ST data (presented in Supplemental Table 2). We employed three complementary metrics to assess spatial structures from distinct perspectives: (1) Moran’s I, which measures spatial autocorrelation, with high values indicating superior spatial continuity; (2) Neighbor MAE, which represents the mean absolute error of the predicted proportions between adjacent spots, where low values denote local smoothness and stability; and (3) Class-neighbor agreement, which quantifies the consistency rate of the predicted classes between adjacent spots to evaluate boundary preservation.

We first assessed spatial consistency on the basis of the controlled synthetic ST data. Here, all the models yielded similar results (Figure 4A). For example, Moran’s I remained within a narrow, robust range of 0.662 to 0.672, whereas neighbor MAE showed minimal local variability (0.117– 0.120). This uniform performance suggests that the spatial structures learned by the BASE and ablation models (NoCls, ExprIn, and NoAAC) resulted in negligible differences under simulation-based conditions. However, transferring these models to real-world human tissue resulted in obvious disparities (Figure 4B). The BASE model maintained exceptional spatial consistency (Moran’s I=0.71; Neighbor MAE=0.084; Class-neighbor agreement=0.084), but the compared models failed. For instance, the ExprIn model (directly utilizing normalized expression values instead of binarized inputs) displayed weakened spatial continuity (Moran’s I = 0.646), high boundary volatility (Neighbor MAE = 0.100), and a reduced capacity to preserve spatial context (class-neighbor agreement = 0.722). Despite having the highest Moran’s I in the simulation, the NoAAC model displayed the lowest spatial continuity for real tissue (Moran’s I = 0.619) and an elevated local error (Neighbor MAE = 0.107).

**Figure 4.**
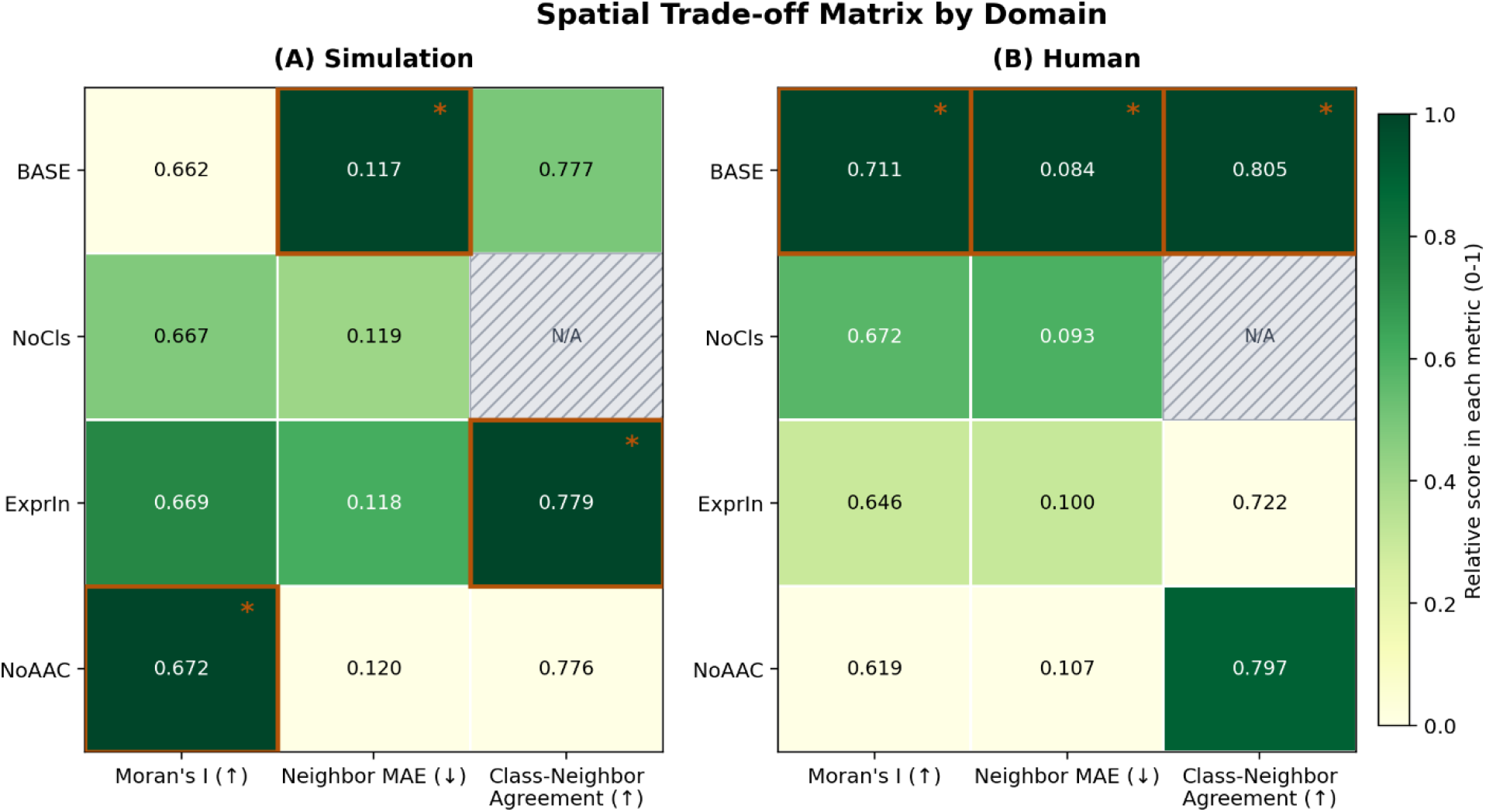
Comparison of spatial consistency between simulation and human domains. The results compare the spatial consistency of each model across (A) the synthetic ST data and (B) real-world human placental ST data. Each row represents a compared model (BASE, NoCls, ExprIn, NoAAC), while each column denotes Moran’s I, neighbor MAE, and class-neighbor agreement. The color of each cell indicates the relative performance across that specific column, illustrating the degree of advantage (with low values being favorable depending on the metric). Asterisks (^*^) and orange borders highlight the top-performing model for each metric.

These results highlight that models with comparable spatial characteristics based on simulations can diverge drastically when subjected to real-world biological noise. Notably, only the BASE model demonstrated robust domain transferability, maintaining stable spatial autocorrelation (0.66 to 0.71 for Moran’s I) and low local variability (0.12 to 0.08 for neighbor MAE) across both domains. These analyses suggest that the superior calibration consistency and classification confidence of the BASE model (Figure 3) serve as the key mechanisms that prevent spatial prediction collapse (tail risk) in cases with real-world ST data.

#### Calibration, Robustness, and Sensitivity

Next, we evaluated error calibration performance using ground-truth labels and synthetic ST data. The distributions of pECE, pMCE, and cTop1ECE across ten random seeds are shown in Figure 5A. While most of the compared models yielded similar calibration metrics, the BASE model exhibited balanced multidimensional reliability (pECE = 0.018±0.003, pMCE = 0.067±0.014, cTop1ECE = 0.012±0.002). For real ST data for human placental tissue, for which mathematical calibration cannot be directly computed because of the lack of ground truth information, we evaluated spatial consistency as a proxy for structural robustness. The seed-specific distributions of Moran’s I, the neighbor mean absolute error (MAE), and class-neighbor agreement based on real tissue data are shown in Figure 5B. For these real-world ST data, the BASE model demonstrated robustness, achieving superior and consistent central tendencies (Moran’s I = 0.701±0.010, neighbor MAE = 0.087±0.002, class-neighbor agreement = 0.808±0.006). In contrast, the compared ablation models exhibited overall inconsistency (Figure 5B).

**Figure 5.**
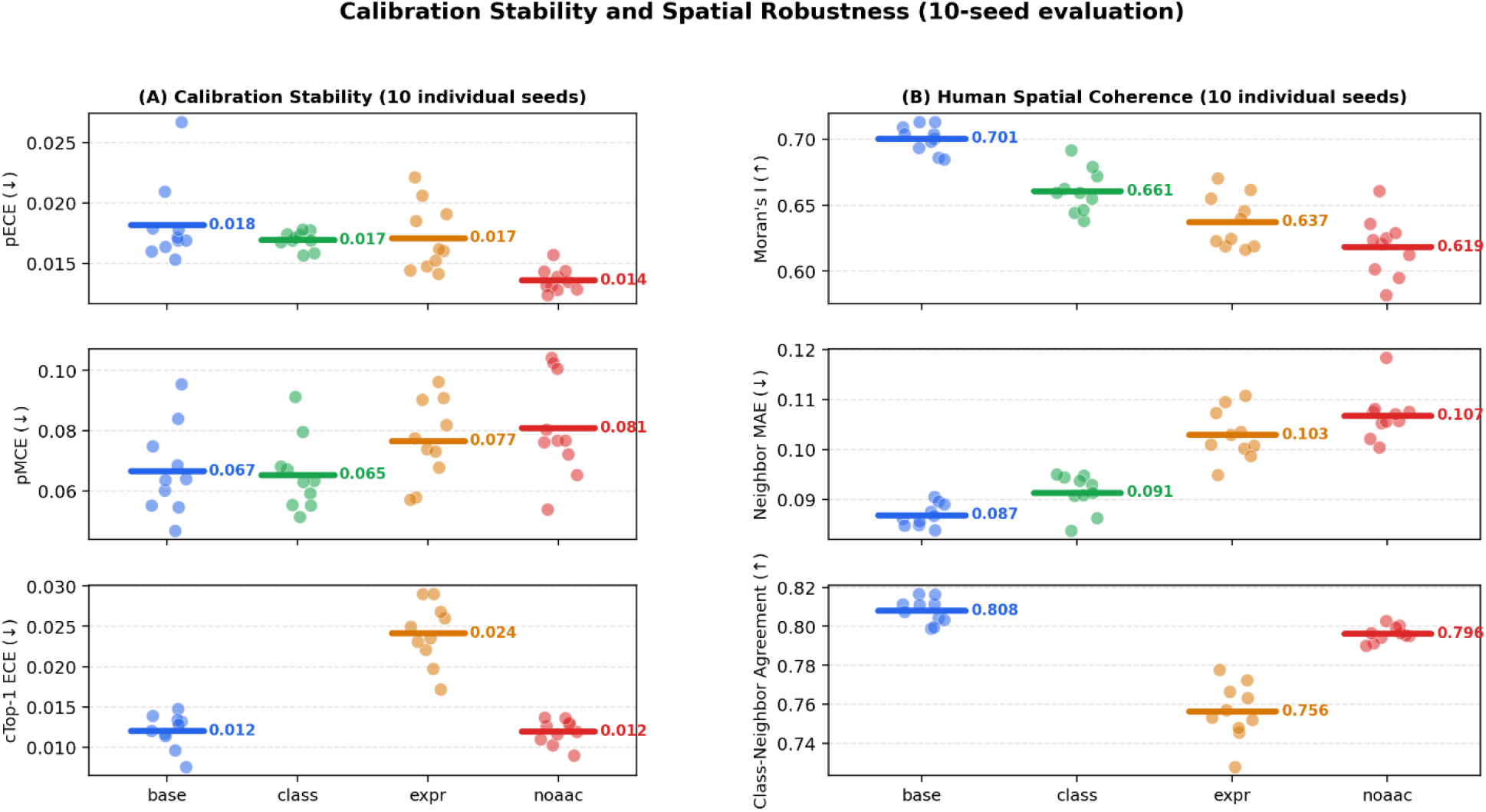
Comparison of calibration stability for the simulation domain and robustness for the real domain across ten random seeds. (A) Error calibration metrics for the synthetic ST data are presented as seed-level strip plots for pECE, pMCE, and cTop1ECE (dots: individual seeds; bold horizontal bars: mean). Low values indicate superior confidence calibration, demonstrating a trade-off relationship in which no single-variant model uniformly dominates all the calibration axes. (B) Using the same seed-level visualization, robustness for the real human tissue ST data and sensitivity to random initialization are evaluated via spatial consistency proxy metrics (Moran’s I, neighbor MAE, and class-neighbor agreement). Across all the weight-initialization seeds, the BASE model exhibits the most stable and reproducible spatial profile, demonstrating robust transferability into the real domain beyond the simulation domain.

Several insights can be gained from this analysis of model sensitivity. First, the average differences in real-world performance between the BASE model and the compared models clearly exceeded their seed-specific standard deviations. Second, this finding indicates that the performance regression observed in the compared models during data transfer stems from reproducible, structural vulnerabilities—dictated by the presence or absence of core features of the model, rather than stochastic sensitivity from random initialization.

Taken together, these results demonstrate that model selection should not rely solely on optimizing a single average metric based on simulations. Securing genuine generalizability requires a dual focus: calibration stability to mitigate worst-case scenarios and reproducible robustness to withstand real-world domain transfer. Under these comprehensive criteria, the full BASE model of GOLDEN successfully avoids overfitting to simulation-specific metrics, achieving the most stable and robust convergence when applied to actual ST data.

### 3.3. Computational requirements of GOLDEN

Beyond predictive accuracy and robustness, the practical adoption of a computational model also depends on its resource requirements and scalability. We therefore examined the computational feasibility of GOLDEN for prospective users by evaluating its parameter scale, memory footprint, and training requirements. GOLDEN features a multitask architecture that integrates a gene-level MLP encoder with a state classification head and a proportion regression head. Accepting the defined 38,584 genes per spot as inputs, the model encompasses approximately 80.16M parameters, which translates to a memory footprint of approximately 305.8 MiB for the weights with FP32 precision. Training was conducted with a single NVIDIA Tesla V100 (32 GB) GPU using a software stack composed of Python 3.10, PyTorch 2.9.1+cu128, and CUDA 12.8. The model was trained with a batch size of 64 over a representative duration of 100 epochs.

## 4. Discussion and conclusion

Overall, the proposed model, GOLDEN, successfully deciphered spatial regions that harbored genetically distinct cells with strong performance and reproducibility. Unlike conventional deconvolution frameworks, which primarily resolve proportions of distinct cell types, GOLDEN addresses genetic heterogeneity, which is a dimension of spatial biology that has remained largely unexplored in this context. GOLDEN effectively mitigates the technical noise inherent in spatial transcriptomics (ST) data through a binarized input representation while seamlessly integrating state classification and proportion estimation via a multitask learning architecture. In tissue microenvironments such as the placenta, where target (fetal) and nontarget (maternal) cells of similar cell types coexist, decoupling the origin proportions remains a notable challenge. GOLDEN addresses this issue by providing a robust computational solution that transcends these data limitations. The robustness of this solution was further confirmed through our ablation study, which revealed that simplified model variants occasionally matched the performance of or slightly outperformed the BASE model in terms of select accuracy metrics but consistently incurred greater worst-case calibration error and yielding weaker confidence alignment, thus underscoring the risks of relying on single average metrics for complex biological data.

The synthetic ST results already underscore the risks of relying on single-metric benchmarks. These performance disparities became even more pronounced upon shifting to data for real-world human placental tissue. While differences in spatial consistency among the variants were negligible during simulation testing, the BASE model demonstrated significantly higher spatial coherence and local structural consistency for actual human data. Crucially, this performance gap was consistently reproduced across ten independent random seeds, which demonstrates that the architecture of GOLDEN avoids overfitting to the training domain, thus ensuring reproducible structural robustness that is resilient to domain shifts.

Beyond the technical performance of GOLDEN, its ability to resolve spot-level genetic origins without relying on external reference profiles is of broader translational relevance. By successfully mapping target and nontarget cell-proportion dynamics with exceptional spatial stability, GOLDEN can serve as a pivotal computational method in the ST domain, with potential value for elucidating complex pathological mechanisms associated with immune responses, such as preeclampsia, allograft rejection, and diverse nonclonal expansion, and for identifying noninvasive diagnostic strategies and niche-specific therapeutic targets in settings such as prenatal monitoring, transplant surveillance, and oncology.

Despite these strengths, certain limitations remain. First, GOLDEN currently focuses on major binary classifications of maternal versus fetal origin harboring over 50% genetic similarity (as presented in Table 1). Thus, further validation of the robustness of the performance of GOLDEN using genetically similar cells, including twins, mosaicism, and mouse lines, is needed. Second, as the current framework is tailored to the spot-level resolution of the 10x Visium platform, improvements in scaling and computational efficiency are necessary to adapt to emerging subcellular-resolution ST technologies, such as Stereo-seq and Xenium.Looking forward, GOLDEN could be extended to accommodate more than two genetic origins by predicting a proportion vector across multiple genotypes rather than a single target fraction. This extension could enable application to settings such as dizygotic twin pregnancies, organ transplantation, or mosaicism, which cannot be represented by the current binary target/nontarget framework.

In conclusion, to the best of our knowledge, GOLDEN is the first approach that can resolve genetic heterogeneity from transcriptional profiles at the spot level.

## Author contributions

Sangjeong Lee: Conceptualization, Algorithm development, and Draft writing; Tae Lyun Ko: Synthetic data generation code development and Draft writing support; Hyoung-Sam Heo: Funding acquisition; Junguee Lee: Pathological annotation; Sung Ki Lee: Funding acquisition and Clinical sample preparation; Hyojung Paik: Conceptualization, Funding acquisition, Formal analysis, Data acquisition, and Draft writing.

## Acknowledgments

This work was also supported by the Korea Bio Data Station (K-BDS), which provided computing resources and technical support.

