## Supplemental data for "GOLDEN: Gene expression and allele-informed decomposition for genetic heterogeneity in spatial transcriptomic data"

Short title: Resolving genetic heterogeneity within a tissue space

**Supplemental Table 1. Real-world ST datasets (Visium) used for evaluation**

| **Sample ID** | **Tissue** | **Preservation** | **Condition** | **Gestational weeks** | **Offspring Sex** | **Median Genes per Spot** | **Spots** | **Note** |
| --- | --- | --- | --- | --- | --- | --- | --- | --- |
| Norm1FxFVisGeni | Placenta | FxF (Fixed Frozen) | Normal | 39 GW | Female | 2,708 | 4,790 | Internal data  GSE xxx  (matched genetic evidence) |
| Pept2FFVis | Placenta | FF (Fresh Frozen) | Preeclampsia | 33+6 GW | Female | 1,568 | 4,357 |  |
| Pla_HDBR9518710  (#PlaHBDR) | Decidua basalis | FF | Normal | 6 PCW | N/A | 3,662 | 1,216 | External data  E-MTAB-12698  (no genetic evidence) |
| WS_PLA_S9101764  (#WSPLAS4) | Decidua basalis, Endometrium | FF | Normal | 8–9 PCW | N/A | 2,970 | 3,568 |  |
| WS_PLA_S9101765  (#WSPLAS5) |  | FF | Normal | 8–9 PCW | N/A | 2,645 | 3,855 |  |
| WS_PLA_S9101766  (#WSPLAS6) |  | FF | Normal | 8–9 PCW | N/A | 1,250 | 3,879 |  |
| WS_PLA_S9101767  (#WSPLAS7) |  | FF | Normal | 8–9 PCW | N/A | 2,707 | 4,186 |  |
| Pla_Camb9518737  (#PlaCamb) |  | FF | Normal | 8–9 PCW | N/A | 2,197 | 4,071 |  |
| WS_PLA_S9101769  (#WSPLAS9) |  | FF | Normal | 8 PCW | N/A | 2,609 | 3,130 |  |
| WS_PLA_S9101770  (#WSPLAS0) |  | FF | Normal | 8 PCW | N/A | 2,436 | 3,355 |  |

Comprising information from two in-house slides and eight slides from a published cohort (Arutyunyan et al., 2023; E-MTAB-12698). Gestational age is reported in gestational weeks (GW; from the last menstrual period) for in-house samples and in postconception weeks (PCW) for the published cohort, reflecting the different sampling stages (late-gestation delivery vs. first trimester). N/A, not applicable.

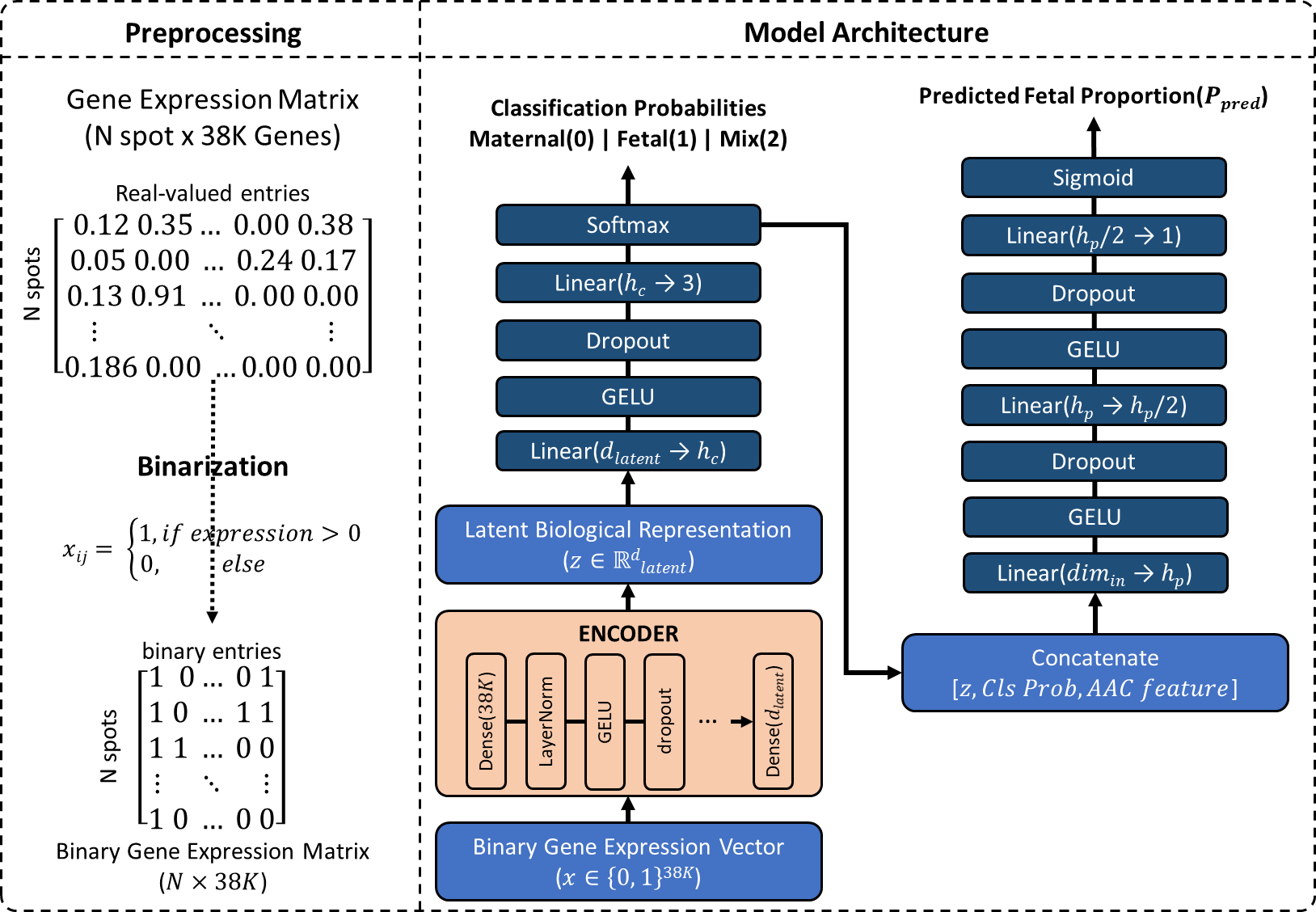

**Supplementary Figure 1.** An overview of the GOLDEN framework | The diagram illustrates the step-by-step pipeline of GOLDEN, from spatial transcriptomic data preparation to multitask prediction. (Preprocessing) The raw gene expression matrix is transformed into a binary gene expression matrix through a binarization process to focus on binary detection patterns. (Model Architecture) For each spot, the binary gene expression vector is processed through an encoder network to generate a low-dimensional latent representation. The classification head uses this latent representation to predict the discrete state of the microenvironment (target, nontarget, or mixed). Finally, the proportion head integrates the predicted state information, the latent representation, and the allele-informed features (AAC) to estimate the continuous target cell fraction.

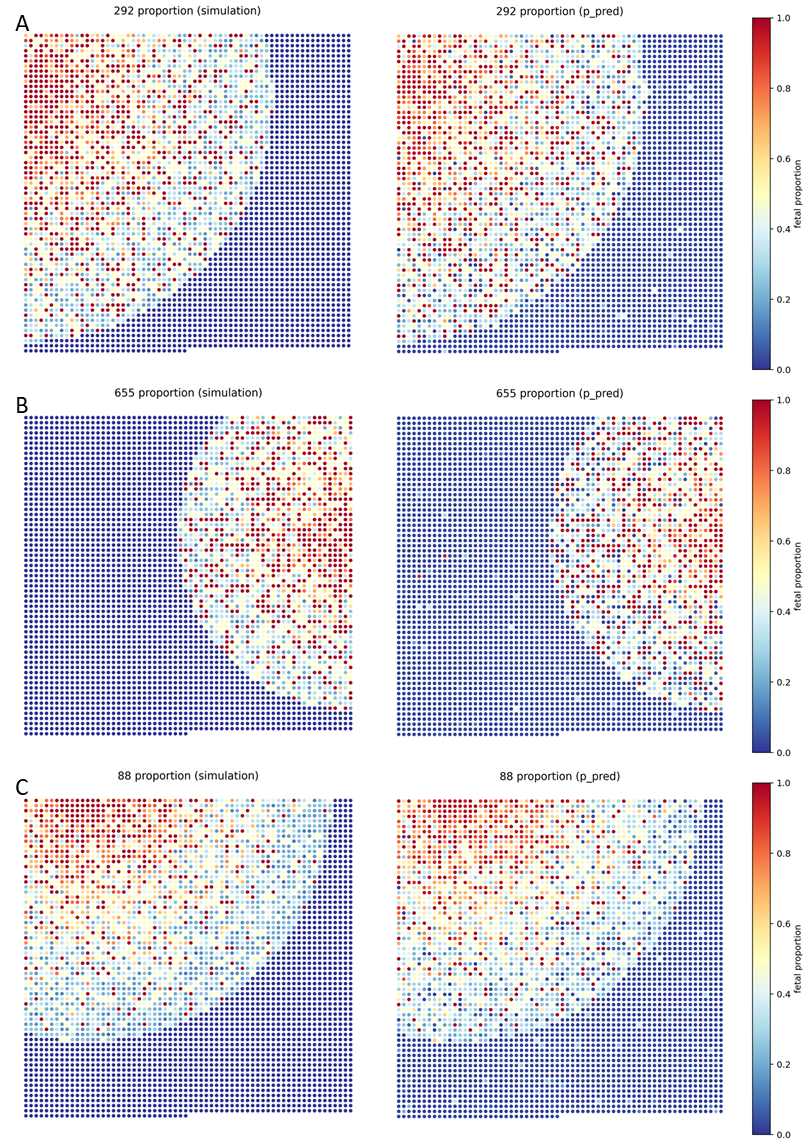

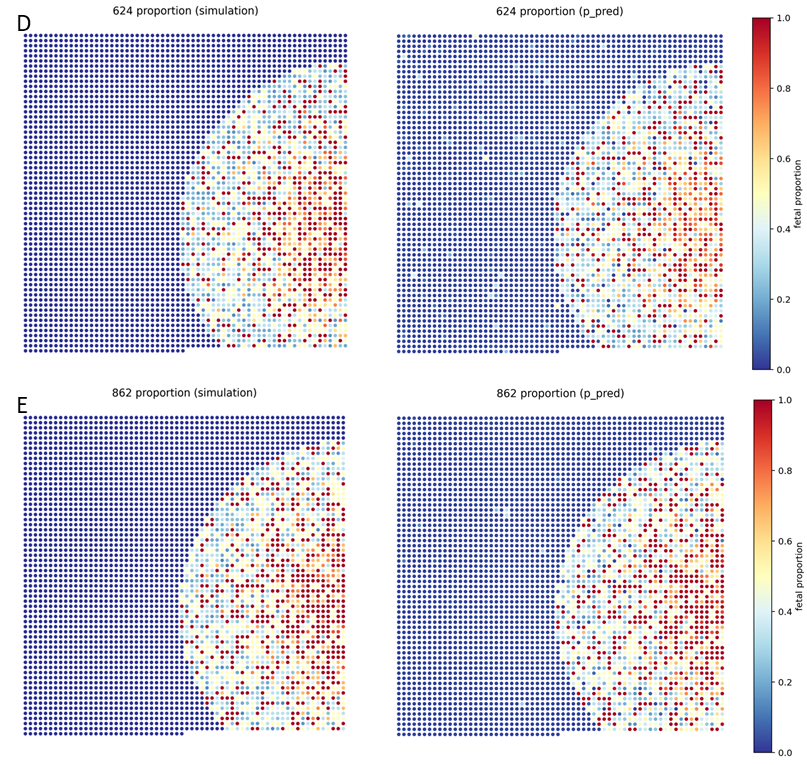

**Supplementary Figure 2.** Spatial distribution comparisons between ground-truth and predicted target cell fractions across six simulation datasets | The spatial plots illustrate a spot-by-spot comparison between the actual ground-truth cell fractions (left) and the corresponding predictions inferred by GOLDEN (right) across all six independent simulation test samples (Simulations 1 to 6). Spatial Reconstruction: For each sample pair, the spatial layout reconstructed with GOLDEN exhibits high visual and spatial alignment with the ground-truth distribution. Sample-Specific Performance Metrics: (A) Simulation 292: MAE = 0.052, RMSE = 0.091, Corr. = 0.970. (B) Simulation 655: MAE = 0.054, RMSE = 0.099, Corr. = 0.962. (C) Simulation 88: MAE = 0.052, RMSE = 0.077, Corr. = 0.972. (D) Simulation 624: MAE = 0.046, RMSE = 0.075, Corr. = 0.976. (E) Simulation 862: MAE = 0.033, RMSE = 0.051, Corr. = 0.990. Across all six simulation datasets, GOLDEN yielded overall average metric values of MAE = 0.045, RMSE= 0.075, and Corr. = 0.977, thus validating the robust generalization ability of GOLDEN across varying spatial mixtures and patterns.

#Placamb

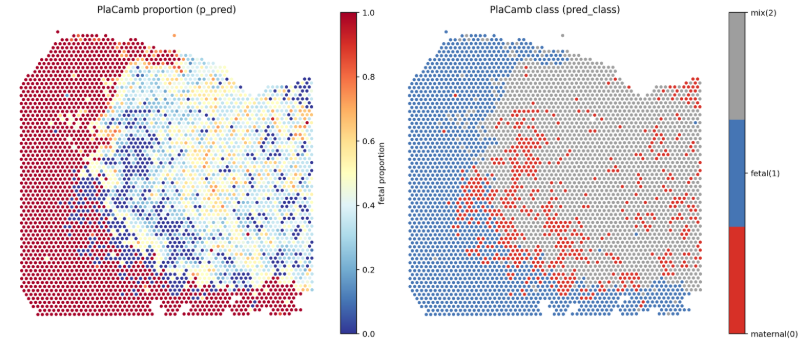

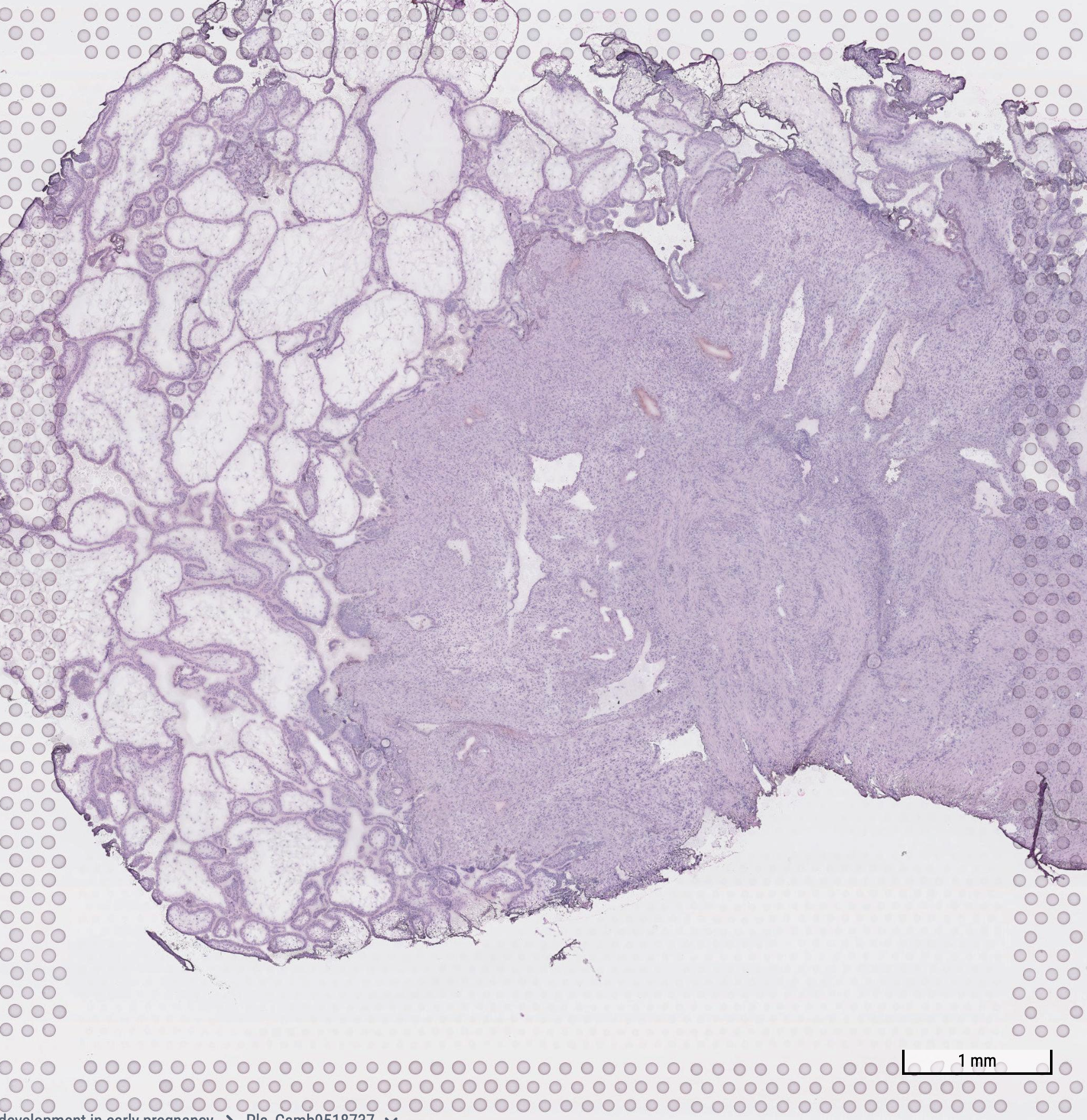

Fetal region

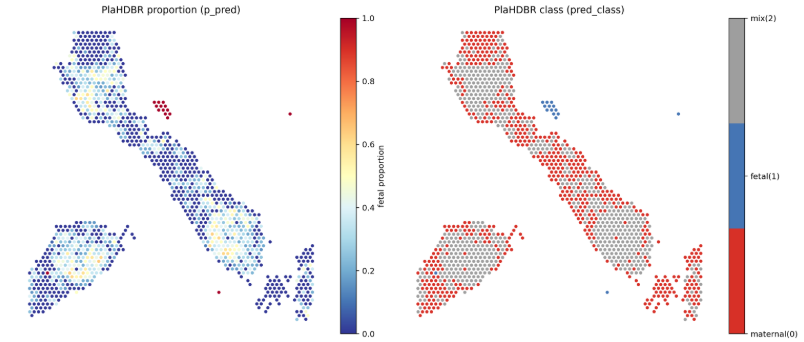

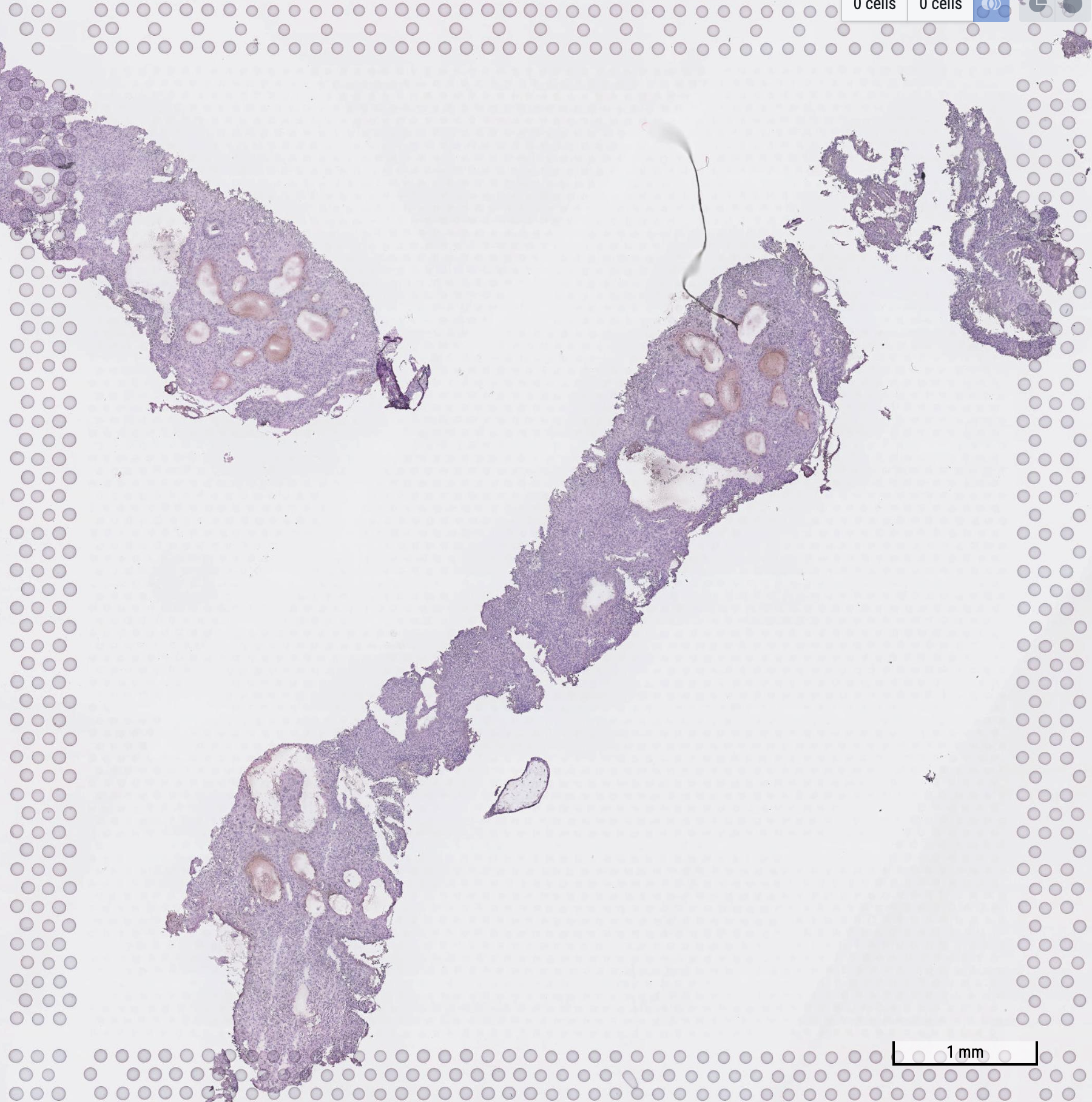

Decidua (mainly maternal)

#PlaHDBR

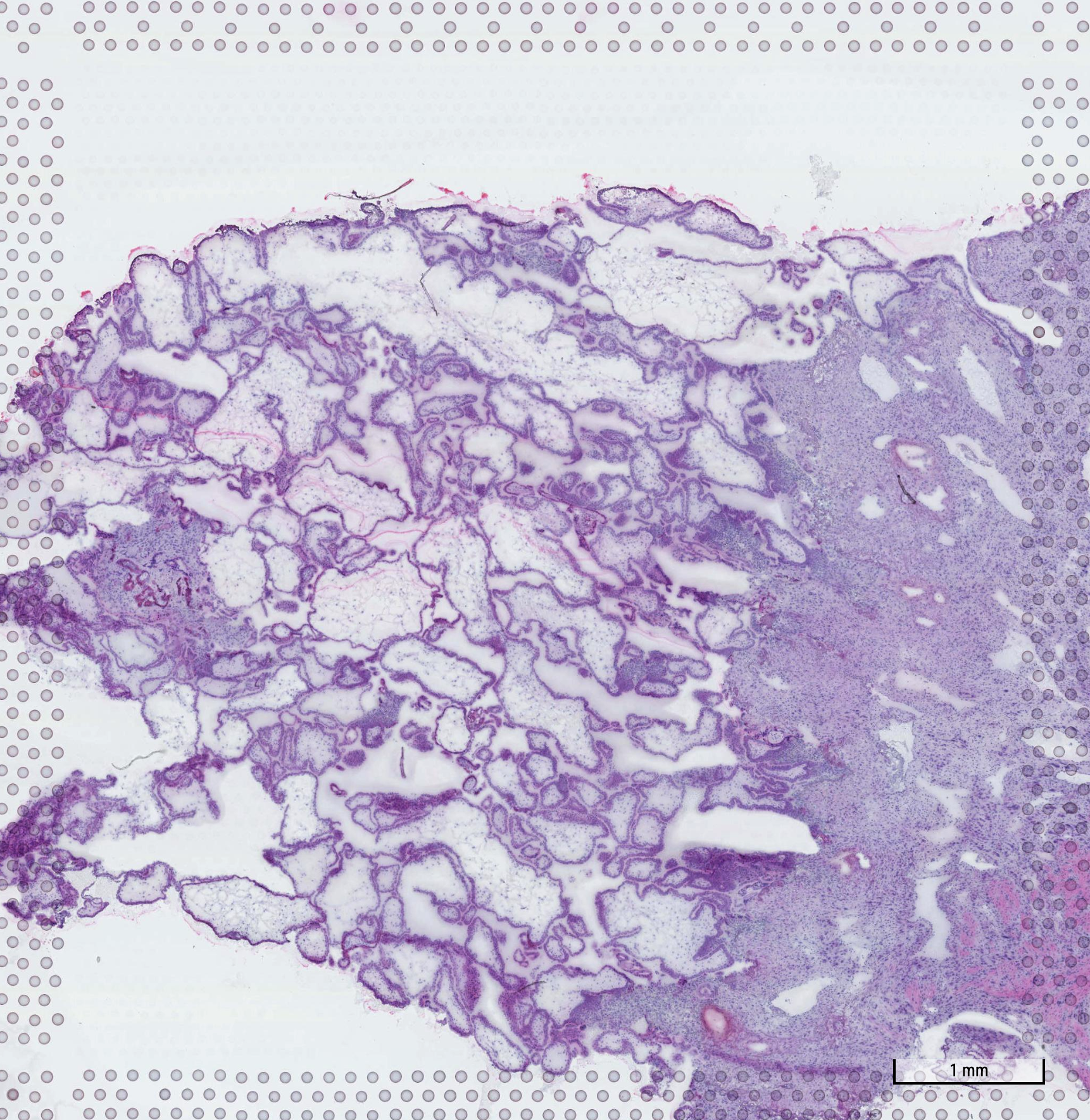

Fetal region

#WSPLAS6

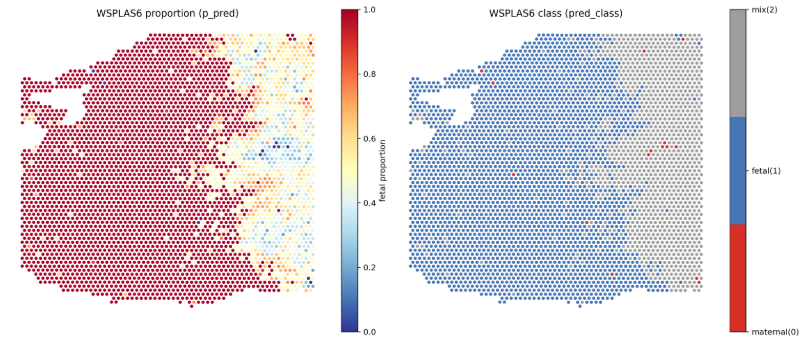

Prediction of GOLDEN

Pathological annotation

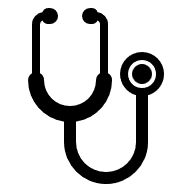

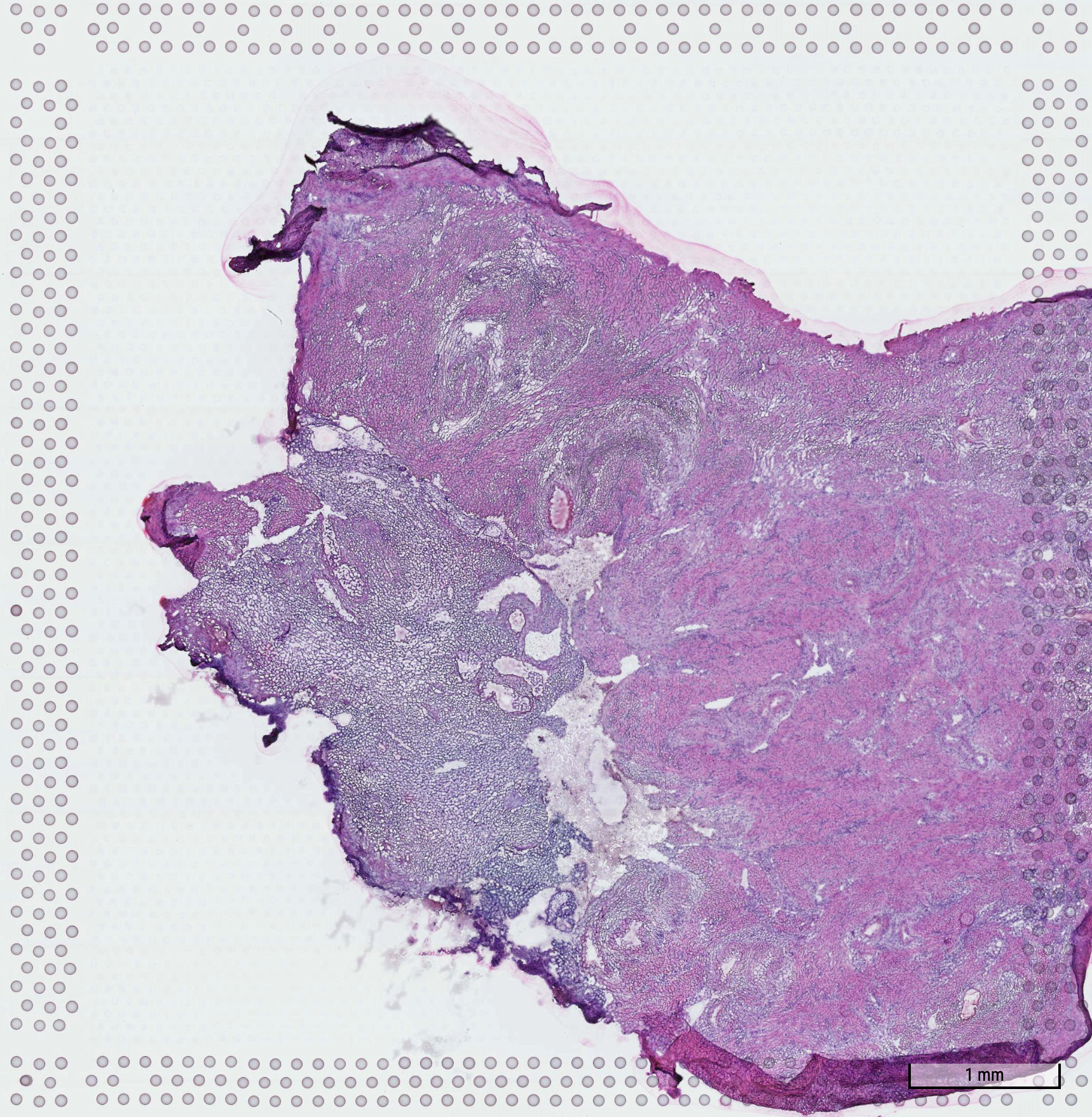

endometrium

(mainly fetus)

uterus

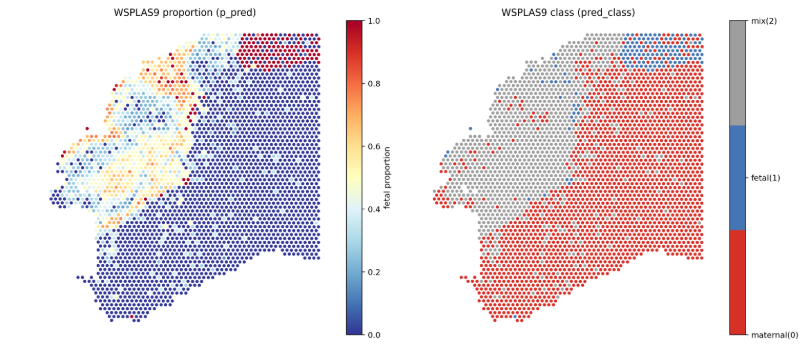

#WSPLAS9

**Supplementary Figure 3.** Comparisons of the spatial distributions of pathological annotations and the predicted target cell fractions for ST data from the external dataset presented in Supplemental Table 1.
